# Dairy products as sources of methanogens for humans

**DOI:** 10.1101/2021.11.16.468822

**Authors:** Cheick Oumar Guindo, Michel Drancourt, Ghiles Grine

## Abstract

Methanogens are detected in human gut from the first moments of life and there is a diversification of methanogens during infancy. However, the sources of acquisition of methanogens are not well elucidated. We therefore investigated 56 dairy products as potential sources of methanogens by applying molecular biology. In the presence of negative controls, we obtained an overall prevalence of methanogens in 85.7% (48/56) of samples by real-time PCR. Further PCR-sequencing identified 73.2% (41/56) of *Methanobrevibacter smithii*. We also found for the first time in dairy products 1.8% (1/56) of *Methanobrevibacter oralis*, 7.1% (4/56) of *Methanobrevibacter millerae*, 1.8% (1/56) of *Methanobrevibacter ruminantium*, 1.8% (1/56) of *Methanocorpusculum* sp. We observed a significant presence (p-value=0.001) of methanogens in fermented dairy products compared to unfermented dairy products. This study gives credit to the fact that dairy products could be considered as a source of methanogens for humans, especially for children.

## 1. Introduction

Methanogens represent the archaea most present in the mammalian microbiota, especially in the human digestive microbiota where they account for 10% digestive tract of the anaerobic microorganisms (Dridi et al., 2011; Guindo, 2020; Nkamga et al., 2017). Methanogens are detected in humans from birth (Grine et al., 2017) and there is a diversification of methanogens in humans over the years: while only *Methanobrevibacter smithii* (*M. smithii*) has been detected and cultured in the neonates (Grine et al., 2017; Koenig et al., 2011; Mihajlovski et al., 2010; Odamaki et al., 2016; Palmer et al., 2007; Sereme et al., 2021), *M. smithii, Methanosphaera stadtmanae* (*M. stadtmanae*), *Methanobrevibacter millerae* (*M. millerae*), *Methanomassiliicoccus luminiyensis, Methanobrevibacter arboriphilicus, Methanobrevibacter oralis* (*M. oralis*), *Candidatus* Methanomethylophilus alvus *and Candidatus* Methanomassiliicoccus intestinalis have been isolated from adult stools (Dridi et al., 2012; Nkamga et al., 2017; Sogodogo, Drancourt, et al., 2019). The various sources for each one of these different species remain unknown. Accordingly, a recent study demonstrated that the presence of methanogens, especially *M. smithii* in children stools is linked to the consumption of dairy products (van de Pol et al., 2017); but this study targeted only two of the methanogens strains present in the human digestive tract, namely *M. smithii* and *M. stadtmanae*. Therefore, we further explored the presence of methanogens in dairy products using molecular biology to target all methanogens currently known in the human digestive tract.

## 2. Materials and methods

### 2.1 Sampling of dairy products

We have investigated the presence of methanogens in different types of dairy products including unfermented dairy products (formula milk, fresh milk, and fresh cheese) and fermented dairy products (yogurt and fermented milk) (Table 1). All these dairy products were purchased in randomly selected supermarkets in Marseille, France in November 2019.

**Table 1.**
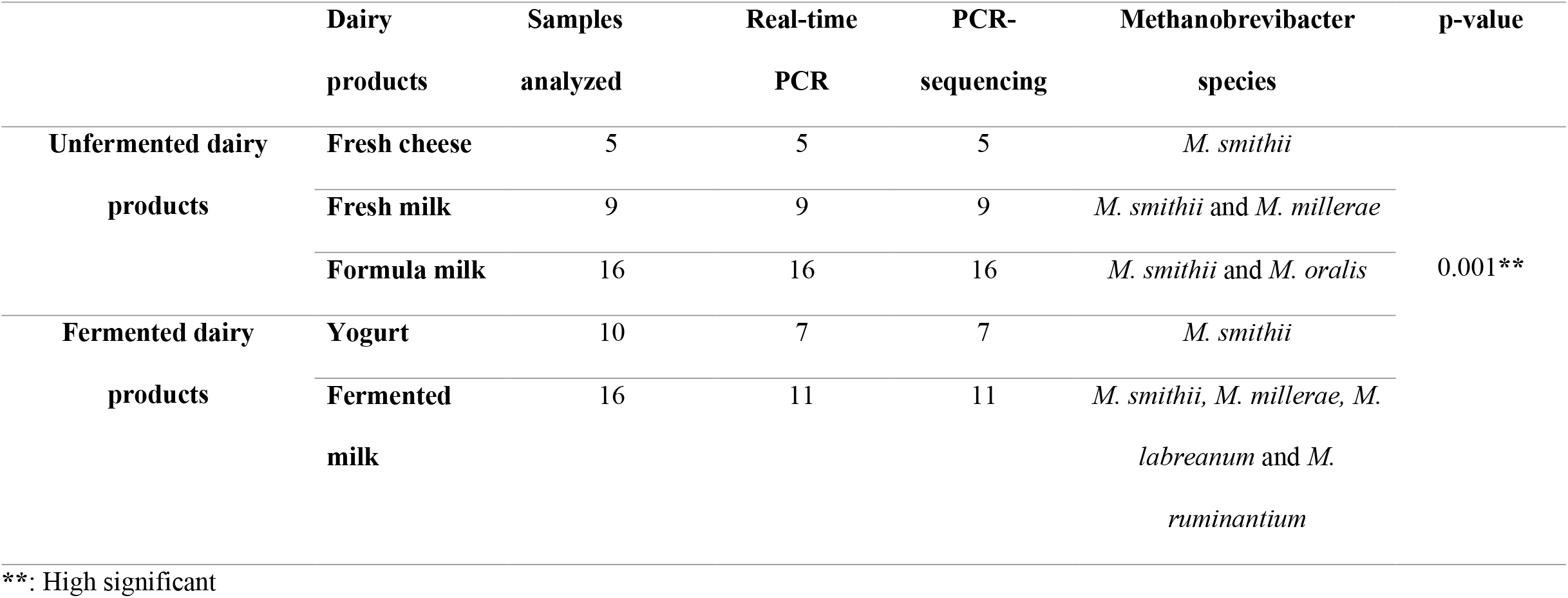
Distribution of dairy products according to the number and results of real-time PCR and PCR-sequencing.

### 2.2 DNA extraction and PCR assays

DNA extraction was performed as previously described (Guindo et al., 2020). Briefly, for cheeses, yogurt and formula milk, 0.2 g was suspended in 200 μL of ultrapure water (Fisher Scientific, Illkirch, France), and a sonication step was performed for 30 minutes. DNA was then extracted with the EZ1 Advanced XL Extraction Kit (QIAGEN, Hilden, Germany) using 200 μL as sample volume and 50 μL as the elution volume. For fresh milk and fermented milk, 200 μL were taken and a sonication step was performed for 30 minutes as above. DNA was then extracted with the EZ1 Advanced XL Extraction Kit (QIAGEN) using 200 μL as the sample volume and 50 μL as the elution volume. The PCR assays targeting the 16S rRNA gene of methanogens, including real-time PCR and PCR-sequencing were performed to investigate the presence of methanogens in dairy products using primer pairs and PCR conditions described previously (Guindo et al., 2021). Sterile phosphate buffered saline (PBS) (Fisher Scientific) was used as a negative control in each DNA amplifications steps.

### 2.3 Phylogenetic analyses

Sequences were edited using ChromasPro software (ChromasPro 1.7, Technelysium Pty Ltd., Tewantin, Australia) as previously used (Guindo et al., 2021; Sereme et al., 2021; Sogodogo, Doumbo, et al., 2019; Sogodogo, Fellag, et al., 2019). Molecular phylogenetic and evolutionary analyses were conducted in MEGA7 as previously described (Kumar et al., 2016).

### 2.4 Statistical analyses

Data were analyzed with RStudio (https://www.R-project.org/) by Fisher. test (** p < 0.01, * p < 0.05, ns: non-significant). We used the former to compare the proportion of methanogen detection by real-time PCR in fermented dairy products compared to unfermented dairy products.

## 3. Results and discussion

We investigated the presence of methanogens in 56 dairy products including 30 unfermented dairy products, and 26 fermented dairy products, of which 48 were positive (Table 1). The overall prevalence of methanogens in dairy products was 85.7% (48/56) by real-time PCR. The prevalence of methanogens in fermented dairy products was 32.1% (18/56) versus 53.6% (30/56) in unfermented dairy products (p-value=0.001). The results here reported were authentified by the fact that negative controls introduced in all experiments, remained negative. PCR-sequencing yielded *M. smithii* in 41 cases, i.e., 73.2% (41/56) exhibiting a 100% sequence similarity with the reference 16S rRNA gene sequence of *M. smithii* ATCC 35061 (accession NCBI: NR_074235) isolated from human stool. *M. smithii* was therefore the most frequent species in dairy products samples (Fig.1). These results are consistent with the literature where *M. smithii* was the only species identified in dairy products (van de Pol et al., 2017). The high prevalence of *M. smithii* in the human digestive tract (Dridi et al., 2009) would therefore be related to the consumption of dairy products (van de Pol et al., 2017). However, we also found for the first time *M. oralis* in one formula milk and this sequence exhibits 99.82% similarity with the sequence of the reference 16S rRNA gene of *M. oralis* CSUR P5920 (NCBI accession: LR590665.1) isolated from Breast milk of healthy breast-feeding mother; *M. millerae* in three fermented milk and in one fresh milk and these sequences have more than 99% similarity with the sequence of the reference 16S rRNA gene of *M. millerae* strain CY31 (NCBI accession: MH443315.1). These two sequences have already been found in human microbiota (Nkamga et al., 2017; Togo et al., 2019). We also found *Methanobrevibacter ruminantium* (*M. ruminantium*) in one fermented milk exhibiting 99.65% similarity with the sequence of the reference 16S rRNA gene *M. ruminantium* strain Yak M20 (NCBI accession: KP123415.1), but also *Methanocorpusculum* sp. T07 in one fermented milk which has 92.24% similarity with the sequence of the reference 16S rRNA gene of *Methanocorpusculum* sp. T07 (NCBI accession: AB288279.1). These two species are only known from the digestive tract of animals especially in the feces of domestic animals (Guindo et al., 2021). In addition, statistical analysis has shown that there is significant correlation between the presence of methanogens in fermented dairy products compared to unfermented dairy products (p-value=0.001), suggesting that fermentation processes might have impact on the DNA of methanogens present in dairy products. This study gives an insight on the concentration of methanogens present in dairy products with the demonstration for the first time ever of the presence of *M. oralis, M. millerae, M. ruminantium* and *Methanocorpusculum* sp. in dairy products. We therefore detected many more species of methanogens compared to the previous study (van de Pol et al., 2017). The presence of *M. smithii, M. oralis* and *M. millerae* in dairy products suggests another mode of acquisition of these methanogens in children through artificial breastfeeding. In addition, a previous study demonstrated that *M. smithii* and *M. oralis* can be acquired in children through breastfeeding (Togo et al., 2019), which strengthens our hypothesis that methanogens present in the digestive tract can be found in milk through mechanisms still unknown. These data add to previous reports of the detection of methanogens in dairy products (van de Pol et al., 2017) and thus giving credit that dairy products could be considered as a source of methanogens for humans, especially for children.

**Figure 1.**
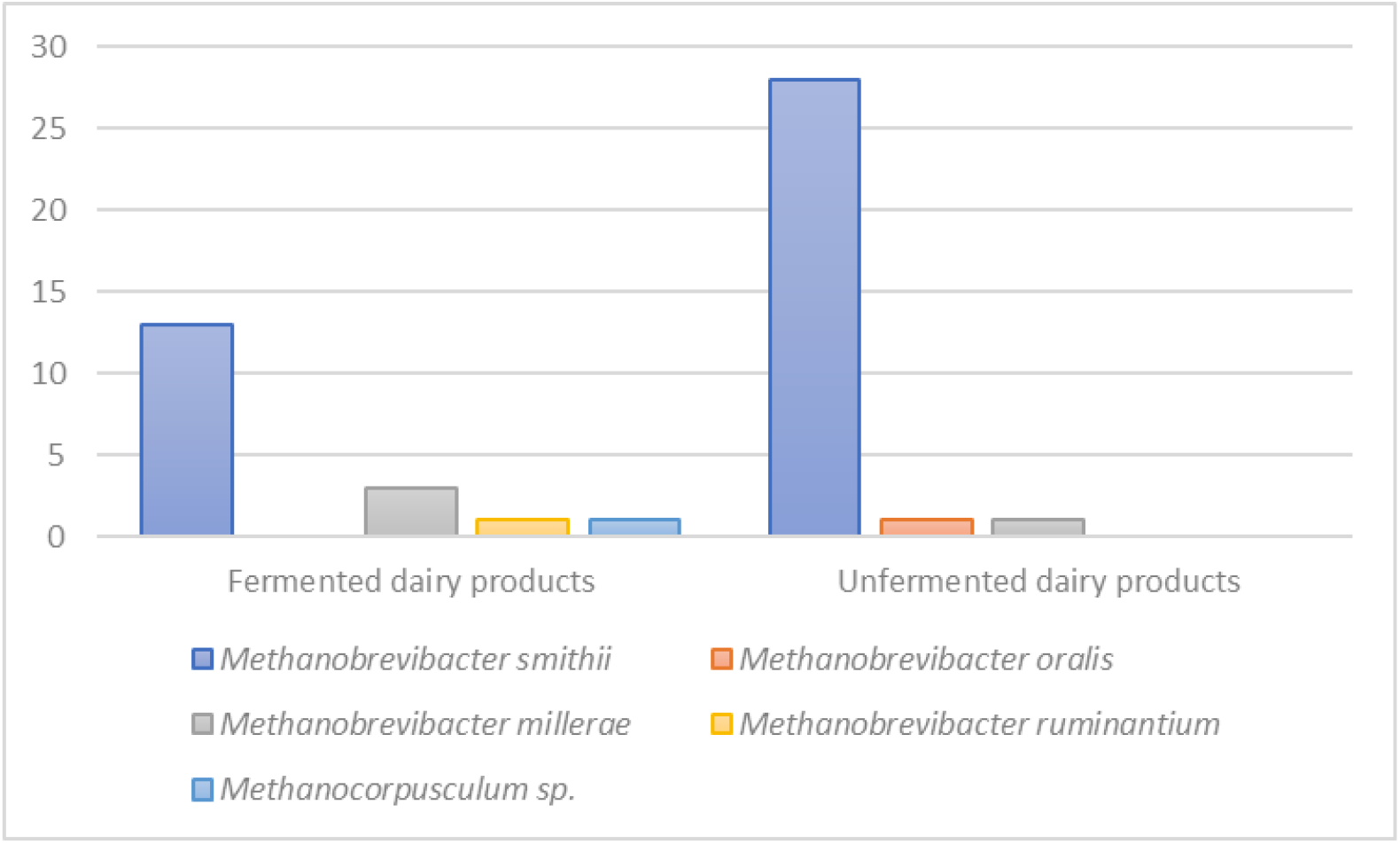
Methanogen species detected in dairy products.

## 4. Conclusions

This study demonstrates, for the first time, the presence of *M. oralis, M. millerae, M. ruminantium* and *Methanocorpusculum* sp. in dairy products and suggests that dairy products may be essential to seed the infant’s microbiota with these neglected critical commensals from the first hours of life.

## ACKNOWLEDGEMENTS

We thank Lanceï KABA for his help in data analysis.

## AUTHORS CONTRIBUTIONS

COG contributed to the collection of samples, conducted the experiments, analyzed the data, and wrote the paper; MD designed the project, participated in the writing of the paper, and provided great support carrying out the experiments; GG designed the project, helped conduct the experiments and participated in the writing of the paper.

